# Discovery of phage determinants that confer sensitivity to bacterial immune systems

**DOI:** 10.1101/2022.08.27.505566

**Authors:** Avigail Stokar-Avihail, Taya Fedorenko, Jeremy Garb, Azita Leavitt, Adi Millman, Gabriela Shulman, Nicole Wojtania, Sarah Melamed, Gil Amitai, Rotem Sorek

## Abstract

Over the past few years, numerous anti-phage defense systems have been discovered in bacteria. While the mechanism of defense for some of these systems is understood, a major unanswered question is how these systems sense phage infection. To systematically address this question, we isolated 192 phage mutants that escape 19 different defense systems. In many cases, these escaper phages were mutated in the gene sensed by the defense system, enabling us to map the phage determinants that confer sensitivity to bacterial immunity. Our data identify specificity determinants of diverse retron systems and reveal phage-encoded triggers for multiple abortive infection systems. We find general themes in phage sensing and demonstrate that mechanistically diverse systems have converged to sense either the core replication machinery of the phage, phage structural components, or host takeover mechanisms. Combining our data with previous findings, we formulate key principles on how bacterial immune systems sense phage invaders.

## Introduction

To survive frequent phage infections, bacteria have developed various anti-phage immune systems. The most widespread of these are restriction-modification (RM) and CRISPR-Cas, both of which cleave invader nucleic acids (Millman et al., 2022). However, studies over the past few years have revealed a plethora of additional defense mechanisms commonly employed by bacteria. These include phage restriction by prokaryotic Argonaute proteins (Garb et al., 2021; Koopal et al., 2022; Kuzmenko et al., 2020; Zaremba et al., 2021), production of small molecules that block phage propagation (Bernheim et al., 2021; Kever et al., 2022; Kronheim et al., 2018), depletion of molecules essential for phage replication (Garb et al., 2021; Hsueh et al., 2022; Ofir et al., 2021; Tal et al., 2022), systems that use small molecule signaling to activate immune effectors (Cohen et al., 2019; Ofir et al., 2021; Tal et al., 2021; Whiteley et al., 2019), retrons that involve reverse transcription of non-coding RNAs (Bobonis et al., 2022; Gao et al., 2020; Millman et al., 2020), and more (Doron et al., 2018; Gao et al., 2020; Goldfarb et al., 2015; Johnson et al., 2022; Millman et al., 2022; Ofir et al., 2018; Rousset et al., 2022; Vassallo et al., 2022).

Although our understanding of bacterial immunity has substantially expanded by these recent discoveries, for the vast majority of the defense systems it is still unknown how they recognize invading phages, and this remains a major unanswered question in the field. Previous studies on individual defense systems, have addressed this question by examining phage mutants that escape defense (Bari et al., 2022; Depardieu et al., 2016; Huiting et al., 2022; Millman et al., 2020; Samson et al., 2013a; Tal et al., 2021, 2022; Yasui et al., 2014). Such phages sometimes evade bacterial immunity by mutations in phage components that activate the bacterial defense system; thus escape mutants can generate valuable insights into the mechanism of defense activation (Samson et al., 2013a). For example, mutations in genes coding for phage inhibitors of the bacterial RecBCD complex enable phages to evade Ec48 retron defense, revealing that the trigger for retron immunity is phage inhibition of the host RecBCD complex (Millman et al., 2020). In another study, phages mutated in the gene responsible for silencing bacterial transcription were found to overcome two different defense systems that deplete deoxynucleotides in response to phage, suggesting that the two systems sense the same pathway during phage infection (Tal et al., 2022). In addition, studying mutant phages revealed that the defense systems PrrC and PARIS trigger abortive infection cell death or growth arrest when they sense phage proteins that inhibit bacterial restriction enzymes (Penner et al., 1995; Rousset et al., 2022), and the mechanism of the Lit abortive infection protein was elucidated based on the observation that T4 phages can avoid Lit activation via mutations in the activator *gol* gene (Bingham et al., 2000; Champness and Snyder, 1982; Yu and Snyder, 1994).

With the understanding that studying phage escape mutants can reveal phage sensitivity determinants of bacterial immune systems, we set out to systematically isolate and study phage escapers for over 50 defense systems (Figure 1). We found that the same phage protein can be sensed by different defense systems, and that central components in the phage replication cycle, including host takeover proteins, phage replication machinery and structural phage proteins, are commonly sensed by bacterial defense systems. Our study shows that bacterial immune systems have converged to sense similar phage components as a marker of infection, and lays the foundations for deciphering the mode of activation for numerous bacterial defense systems.

**Figure 1.**
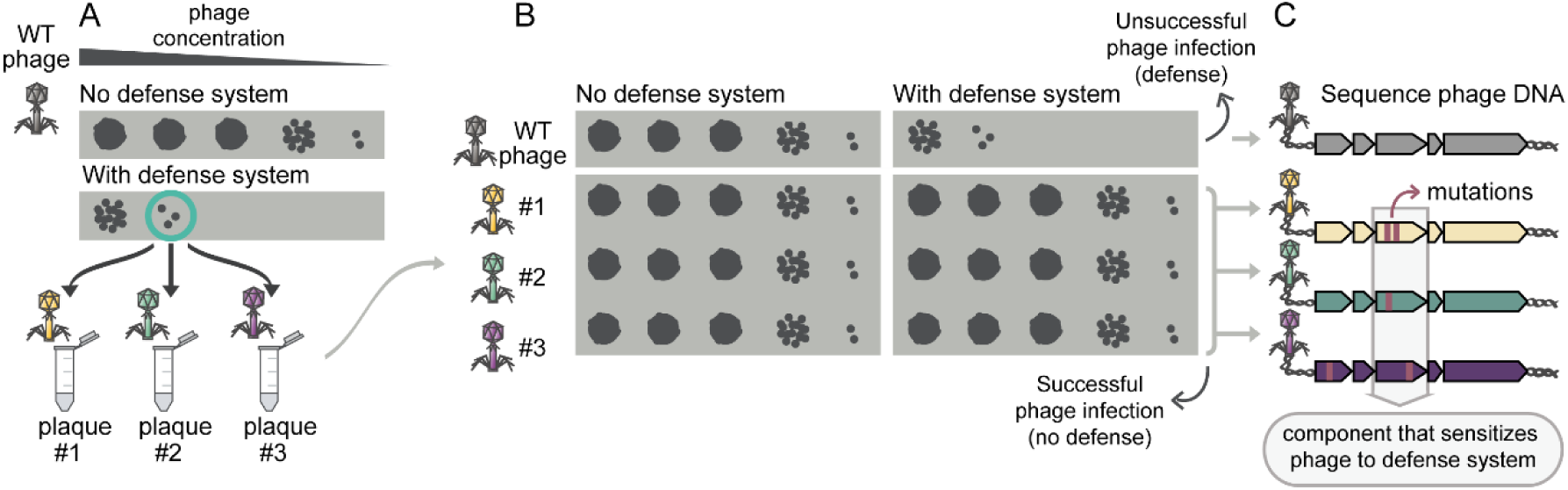
Schematic of the pipeline for isolation and study of phages that overcome defense. (A) Active defense systems cloned into *E. coli* or *B. subtilis* provide protection against phage infection, as observed using serial dilution plaque assays. Individual phage plaques that are formed on bacteria expressing the defense system are collected to screen for escaper phage mutants. (B) The collected plaques are tested for their ability to overcome the defense system using serial dilution plaque assays. A phage is considered to escape defense if it is less sensitive to the defense system than its WT ancestral phage. (C) Phages that escape bacterial defense are subjected to whole genome sequencing. The genome sequences of different isolates are compared to identify loci commonly mutated in different phages, likely explaining their ability to overcome defense.

## Results

We collected a set of recently identified defense systems for which the activating phage trigger is unknown. These included four CBASS systems (Cohen et al., 2019; Hobbs et al., 2022), eight retron systems (Millman et al., 2020), two SIR2-dependent systems (Garb et al., 2021), as well as representatives of the BREX, DISARM, and Viperin systems (Bernheim et al., 2021; Goldfarb et al., 2015; Ofir et al., 2018) and 38 additional systems of mostly unknown function, described in Doron et al. (2018), Gao et al. (2020), and Millman et al. (2022) (Figure 2). Each system was cloned from its source organism and introduced into either *E. coli* or *B. subtilis*, and in some cases we used several instances of the same system from different source organisms. Altogether, the collection of defense system strains examined in the current study included 62 strains representing 54 different defense systems (Figure 2, Table S1).

**Figure 2.**
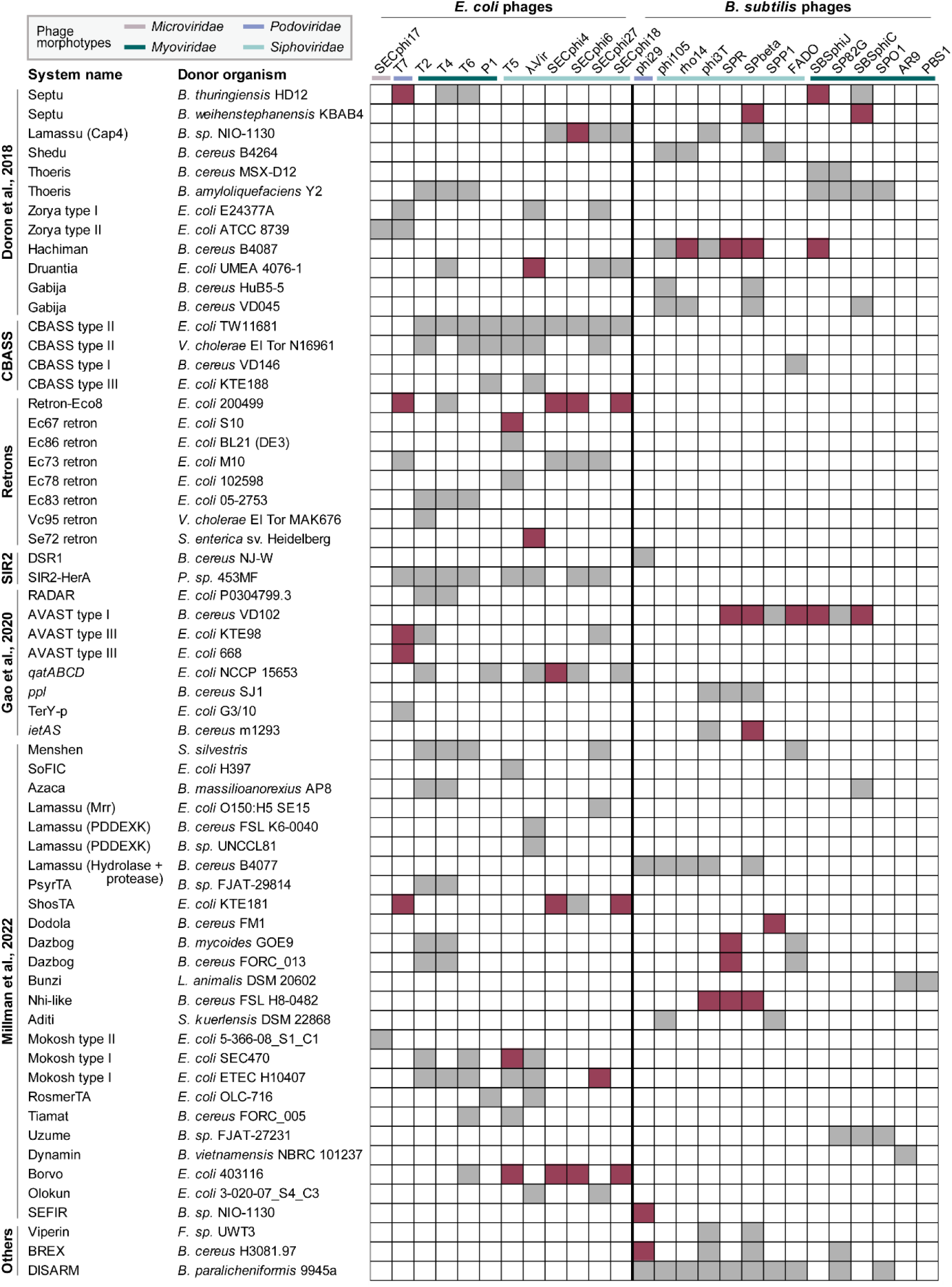
List of defense systems and phages included in screen. Defense systems were expressed, usually under their native promoter, in either *E. coli, B. subtilis* or both, and subjected to an array of phages infecting the relevant host. Cases in which the system protects against a given phage are marked by a filled square. Maroon denotes cases for which at least one escaper phage was isolated, and grey denotes cases for which escaper phages were not isolated. The full names of the donor organisms for each system are provided in Table S1.

Bacteria expressing each of the defense systems were infected with a panel of 12 *E. coli* and 14 *B. subtilis* phages, including representatives from the major phage morphotypes *Myoviridae*, *Siphoviridae*, *Podoviridae*, and *Microviridae* (Figure 2). Despite the efficient protection that these systems provide, in many cases a small number of phage plaques still formed upon infection of bacteria expressing the system. We hypothesized that some of these plaques contain phages with a spontaneous mutation enabling them to escape bacterial defense. These escaper phages were isolated and their ability to escape defense was verified and quantified by serial dilution plaque assays (Figure 1A-B). In most cases, the escaper phages generated more plaques on strains encoding the defense system as compared to WT phages, while in some cases the escape phenotype was manifested in larger plaques of the escaping phages (Figure 3A, Figure S1). Overall, we screened 199 phage-host pairs and successfully isolated 192 phages that partially or fully overcame 19 out of the 54 defense systems included in the study (Figure 3A).

**Figure 3.**
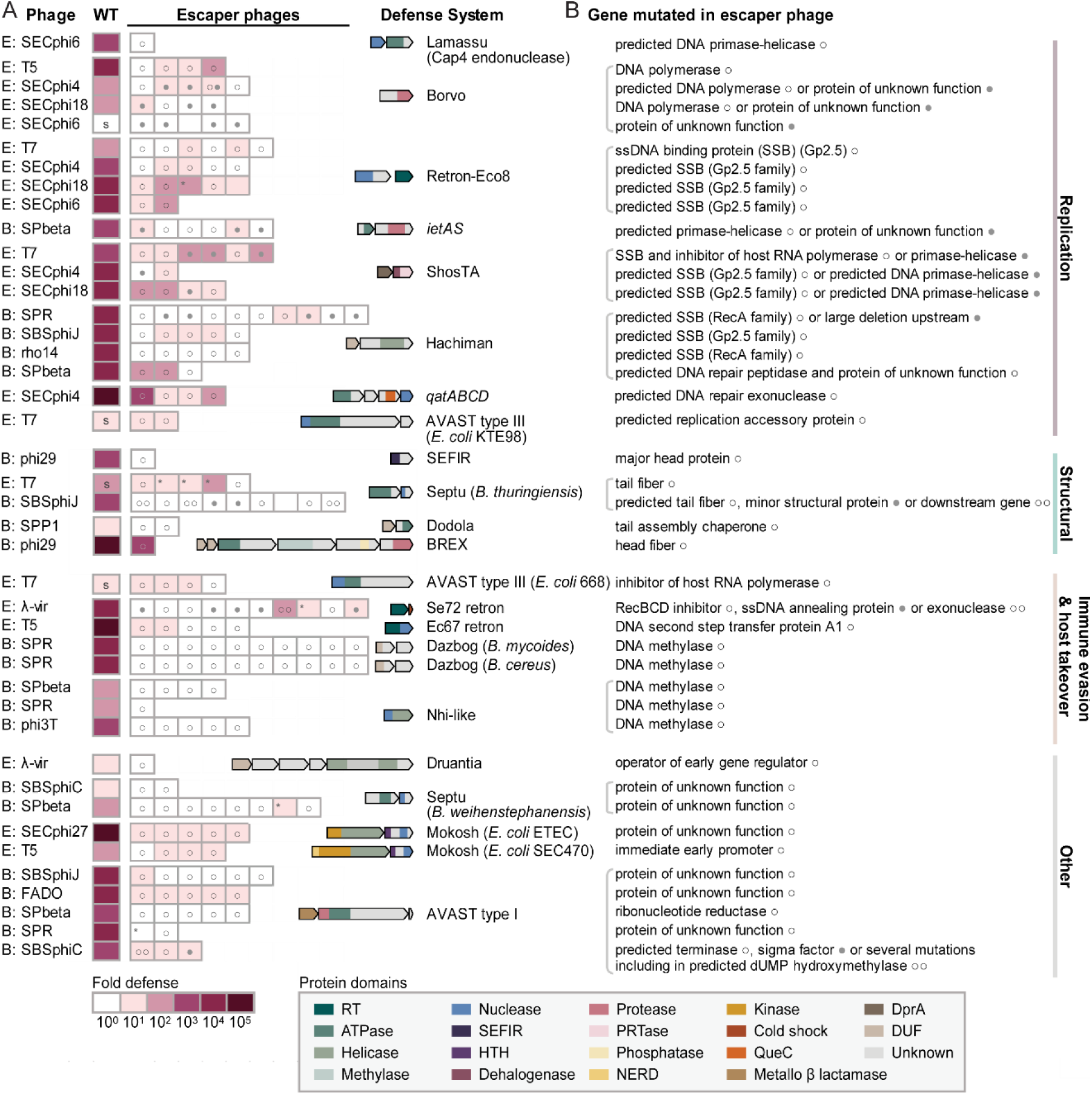
Phages evolve to evade diverse defense systems. (A) Anti-phage systems for which escaper phages were isolated. The isolated phage name is designated to the left, “B” and “E” mark *B. subtilis* and *E. coli* phages, respectively. Fold defense was measured using serial dilution plaque assays and calculated as the ratio between phage PFUs obtained on defense-containing cells and on control cells that lack the defense system (see Figure 1B). Data represent average of two independent replicates. A designation of ‘s’ stands for a marked reduction in plaque size on defense-containing cells compared to control cells. In these cases, escapers showed larger plaques on defense-containing cells (see Figure S1). An Asterisk on the top left of the rectangle indicates cases for which no mutation was detected in the phage genome sequence. Circles (empty or full) are used to specify the gene mutated in each phage isolate as detailed in panel B. (B) The identity of genes mutated in phage isolates that escape the defense activity of each of the anti-phage systems (see mutation details in Table S2).

To identify the possible cause for phage escape from each of the defense systems, we sequenced the full genome of each of the 192 escaper phages and compared them to the sequence of the WT phage, which is sensitive to defense (Figure 1C). In 79% of the cases there was only a single mutation in the escaper phage genome; in cases where escaper phages had mutations in multiple genes, we searched for a gene that is commonly mutated in several different escaper phage isolates (Figure 1C, Figure 3B, Table S2). We found that mutated genes in escaper phages could mostly be categorized as being involved in host takeover and immune evasion, DNA replication, or in building the phage structural components (Figure 3B). Below we focus on several categories of defense systems and the modes by which phages escape them.

### Phage triggers for retron defense systems

Retron defense systems consist of three components: a reverse transcriptase (RT), a non-coding RNA, and an associated effector protein (Millman et al., 2020). The retron RT reverse transcribes the non-coding RNA to form a chimeric RNA-DNA molecule which, together with the RT, has been hypothesized to sense phage infection. Defense by retrons involves cell suicide or growth arrest in response to phage infection (abortive infection), which is carried out by the associated effector protein (Bobonis et al., 2022; Millman et al., 2020). Based on phage escaper data, it was previously shown that retron Ec48 “guards” the cellular RecBCD complex, such that when phages inhibit RecBCD, the retron becomes active and leads the cell to growth arrest (Millman et al., 2020). Studies on another retron, St85, demonstrated that phage proteins that degrade or modify the retron reverse-transcribed DNA activate the toxicity of the retron effector (Bobonis et al., 2022). However, it is unknown how other retrons sense phages.

In the current study, we attempted to isolate defense-insensitive phages for eight retron defense systems, and successfully isolated phage escapers for three of them (Figure 2). Altogether, 33 escaper phages were isolated: 10 for the Se72 retron system, 18 for Retron-Eco8 and 5 for retron Ec67 (Figure 3A, Table S2).

Of the ten λ-vir phages that escaped Se72 retron defense, nine contained mutations in the operon that encodes the RecBCD inhibitor protein Gam. Mutations were found either in *gam* itself or in the two other genes in that operon, *bet* or *exo*, encoding a ssDNA annealing protein and a 5′-3′ exonuclease, respectively (Figure 3B, Table S2). This suggests that a component of the *gam-bet-exo* operon may be the trigger for Se72 defense and activate its toxic effector. To check whether either of the proteins encoded by this operon is sufficient to activate the Se72 retron, we tried to co-express each of the genes together with the defense system and monitored bacterial growth for signs of activation of the toxic effector. Expression of *gam* (but not *bet* or *exo)* was clearly toxic in cells that contain the Se72 retron defense system, but not in bacteria lacking Se72 (Figure 4A). Expression of the mutated version of *gam*, found in escaper phages that overcame Se72 mediated defense, was not toxic in retron-containing cells (Figure 4A). These data indicate that the λ phage Gam protein is likely the phage component that activates Se72 retron defense. We hypothesize that mutations in the *bet* or *exo* genes lead to escape from retron defense because they have a polar effect on the co-transcribed *gam* gene.

**Figure 4.**
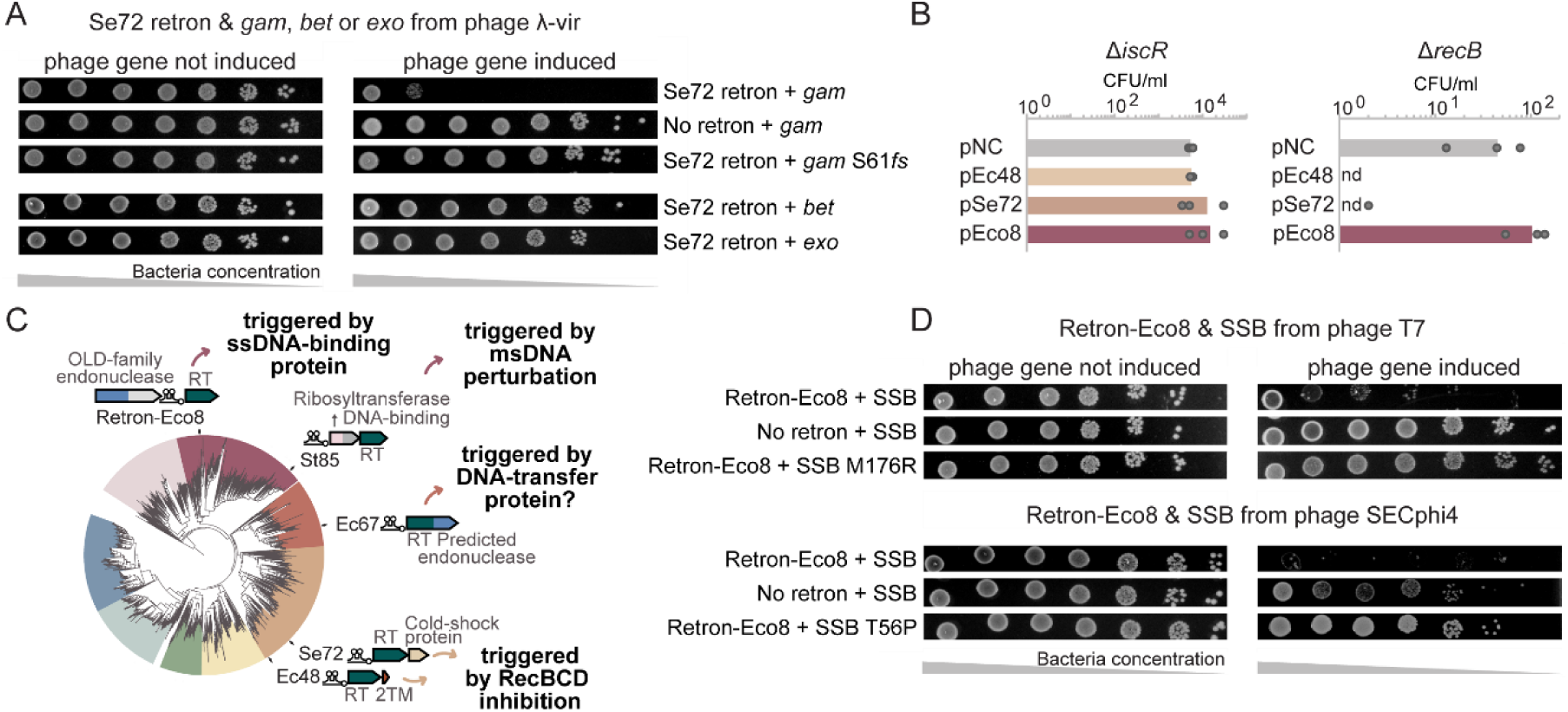
Phage triggers for retron defense systems. (A) Expression of a phage-encoded RecBCD inhibitor activates toxicity in a strain that contains the Se72 retron system. Bacterial viability was measured for cells encoding the Se72 defense system and a single phage gene (either *gam, bet* or *exo*) by plating ten-fold serial dilutions of cells in conditions that repress or induce expression of the phage gene. “No retron” is a negative control strain containing an empty vector instead of a retron. *gam*S61fs is a *gam* version with a frameshift mutation, found in phages that overcome Se72 (Table S2). (B) A vector that contains the Se72 retron anti-phage system cannot be transformed into cells lacking RecB. Transformation efficiency of a plasmid containing either the Ec48, Se72 or Eco8 anti-phage retron systems or an empty vector as a negative control (pNC) into an *E. coli* strain with a deletion of either *recB* or *iscR*. The number of colony-forming units (CFU) per milliliter following transformation is presented. Bar graphs represent an average of three replicates, with individual data points overlaid. “nd”, not detected, indicates cases for which no colonies were detected (limit of detection = 1 CFU/ml). (C) Phylogenetic analysis of ∼4,800 homologs of retron reverse transcriptases, adapted from (Millman et al., 2020). Selected retrons from the tree are marked, with the predicted phage-encoded activator denoted. The triggering mechanism for the St85 retron is reported based on a previous study (Bobonis et al., 2022). (D) Expression of a phage-encoded ssDNA binding protein (SSB) activates toxicity in a strain that contains the retron Eco8 system. Bacterial viability was measured for cells encoding the Eco8 defense system and the WT and mutated SSB proteins from either phage T7 or SECphi4. The experiment was performed as described for panel A.

As the λ phage Gam protein is an inhibitor of the host RecBCD complex, we hypothesized that, as in the case of the Ec48 retron system (Millman et al., 2020), the Se72 retron may function by monitoring the integrity of the host RecBCD complex. To test whether perturbation of RecBCD triggers the toxic activity of the Se72 retron system, we checked if the Se72 system can be expressed in cells that lack *RecB*. While an empty control vector was transformed with high efficiency into Δ*recB* cells, a vector encoding the Se72 system appeared toxic in Δ*recB* cells, transforming >60-fold less efficiently as compared to the control (Figure 4B). The same Se72-containing vector transformed with high efficiency into a control strain deleted in an unrelated gene (Δ*iscR*), indicating that the perturbation of the RecBCD complex activates the toxicity of the Se72 retron system. Together, our results suggest that retrons Ec48 and Se72 both guard the RecBCD complex. Interestingly, these two systems differ in their operon composition: the Ec48 system encodes an effector protein with predicted transmembrane helices, while the Se72 system harbors an effector with a cold-shock protein domain that has no homology to the effector of Ec48. By contrast, these two retrons share close homology in their RT proteins (Figure 4C, Figure S2C). These data support a model in which the retron RT, possibly in combination with the non-coding RNA/DNA component, is involved in RecBCD sensing.

We have previously shown that Retron-Eco8 readily transforms into cells deleted in *recB*, which indicated that this retron is triggered via a mechanism possibly unrelated to RecBCD inhibition (Millman et al., 2020) (Figure 4B). To investigate the phage trigger for this retron, we examined 18 mutant phages isolated from plaques formed on cells expressing the Retron-Eco8 system. These included six escape mutants of phage T7 and 12 mutants of a group of phages that are closely related to each other; phages SECphi4, SECphi6, and SECphi18 (Figure 3, Table S2). In all T7 escapers we found either one or two missense point mutations in the gene encoding the phage single-stranded DNA binding protein (SSB), which is essential for T7 phage DNA replication (Kim and Richardson, 1993). The T7 SSB interacts with the phage DNA polymerase and primase-helicase proteins to form the replication complex, where it binds and protects ssDNA intermediates during replication (Hollis et al., 2001). Intriguingly, ten of the twelve SECphi4, SECphi6 and SECphi18 phage mutants that escaped Retron-Eco8 were also mutated either in the promotor or coding sequence of a predicted SSB protein that has structural homology with the T7 SSB (Figure S2A), indicating that distantly-related phages can escape Retron-Eco8 defense via mutations in their SSB protein (Figure 3B, Table S2). Co-expression of Retron-Eco8 with the SSB protein of either the T7 or the SECphi4 phages resulted in cellular toxicity, and this toxicity was alleviated when these SSB proteins harbored the point mutations found in the escaper phages (Figure 4D). These results suggest that the phage SSB protein forms the trigger for the abortive infection function of the Retron-Eco8 system. Possibly, triggering may involve interactions between the reverse-transcribed ssDNA formed by the retron and the SSB protein.

Five escape mutants in phage T5 were isolated for the retron Ec67 defense system (Figure 3, Table S2). All five isolates had missense mutations in the T5 phage *A1* gene (either I36T, S47P, A15T, C33R or V214M), and one of them had an additional mutation in the stop codon that extended the A1 protein reading frame by 17aa (Table S2). A1 is an essential early-expressed phage protein required for the full transfer of T5 DNA into the host cell, and A1 mutants display various phenotypes including defects in degradation of host DNA and in host RNA and protein synthesis shut-off (Davison, 2015). Structural modeling of the A1 protein reveals a predicted nuclease domain at the C-terminus and a predicted N-terminal DNA-binding domain (Figure S2B). Four of the mutations found in T5 phages that evade Ec67 retron defense localized to this predicted DNA-binding domain, implying that the DNA-binding activity of the A1 protein may be involved in sensitizing T5 phages to the Ec67 retron defense system. The A1 protein is highly toxic to *E. coli* (Klimuk et al., 2020), and we therefore could not check whether expression of A1 alone activates Ec67 retron toxicity.

A common theme among retron-interacting proteins found in this and other studies, including T7 SSB, T5 A1, Rac prophage DNA exonuclease, *E. coli* DNA methylase (Bobonis et al., 2022) and RecBCD, is that they all have the capacity or predicted capacity to interact with DNA. It is possible that interaction of these proteins with the retron-generated ssDNA forms part of the basis for their sensing by the retron system.

### Identifying new abortive infection systems and their phage triggers

For the vast majority of anti-phage systems, it is still unknown how the system defends against infection. In the case of the defensive retrons, which provide protection by an abortive infection mechanism, our data showed that phages can escape defense via mutations in the phage protein that would normally trigger the system’s toxicity. Expression of this protein alone was sufficient to trigger the toxic activity of the system (Figure 4A,D) (Millman et al., 2020). Based on this, we reasoned that the genes identified in our screen as mutated in escaper phages could be utilized to expose anti-phage systems, other than retrons, that function via abortive infection in response to a specific phage protein. If anti-phage defense functions via abortive infection, co-expression of the defense system with the respective activator phage protein is expected to elicit cellular toxicity.

To identify systems potentially functioning via abortive infection, we paired each system with the phage gene found to be mutated in escaper phages isolated on that system. We then expressed these phage genes in cells that contain the respective defense system, and examined the bacterial colonies for signs of toxicity (Figure S3, Figure 5A-E). Altogether, 28 pairs of defense system/phage gene were tested. In most of the cases, expression of the phage gene was not toxic in cells that contained the defense system (Figure S3A), suggesting that either the system does not elicit cellular toxicity when activated, or that the specific phage gene, when expressed alone, is necessary but not sufficient to activate abortive infection (as recently suggested for Pycsar (Tal et al., 2021) and CBASS (Huiting et al., 2022)). In several other cases, expression of the phage gene exhibited non-specific toxicity in both defense system-containing cells and in control cells lacking the system (Figure S3B). In five cases, expression of the phage gene was toxic in defense system-containing cells but not in control cells that lack the system, revealing a suspected abortive infection function for the systems Hachiman (Doron et al., 2018), *ietAS* (Gao et al., 2020), Borvo, Mokosh type I, and Dazbog (Millman et al., 2022) (Figure 5A-E). Co-expression of the defense system with the mutated version of the phage gene alleviated toxicity either fully or partially, explaining how mutant phages escaped defense (Figure 5A-E). Infection experiments in liquid cultures further supported the hypothesis that these systems protect via abortive infection, since introduction of phage at a high multiplicity of infection (MOI) resulted in culture collapse or growth arrest, while the cultures survived infection at low MOI (Figure 5F) (Lopatina et al., 2020). These results reveal an abortive infection phenotype, as well as the phage trigger, for systems whose mode of action was so far unknown.

**Figure 5.**
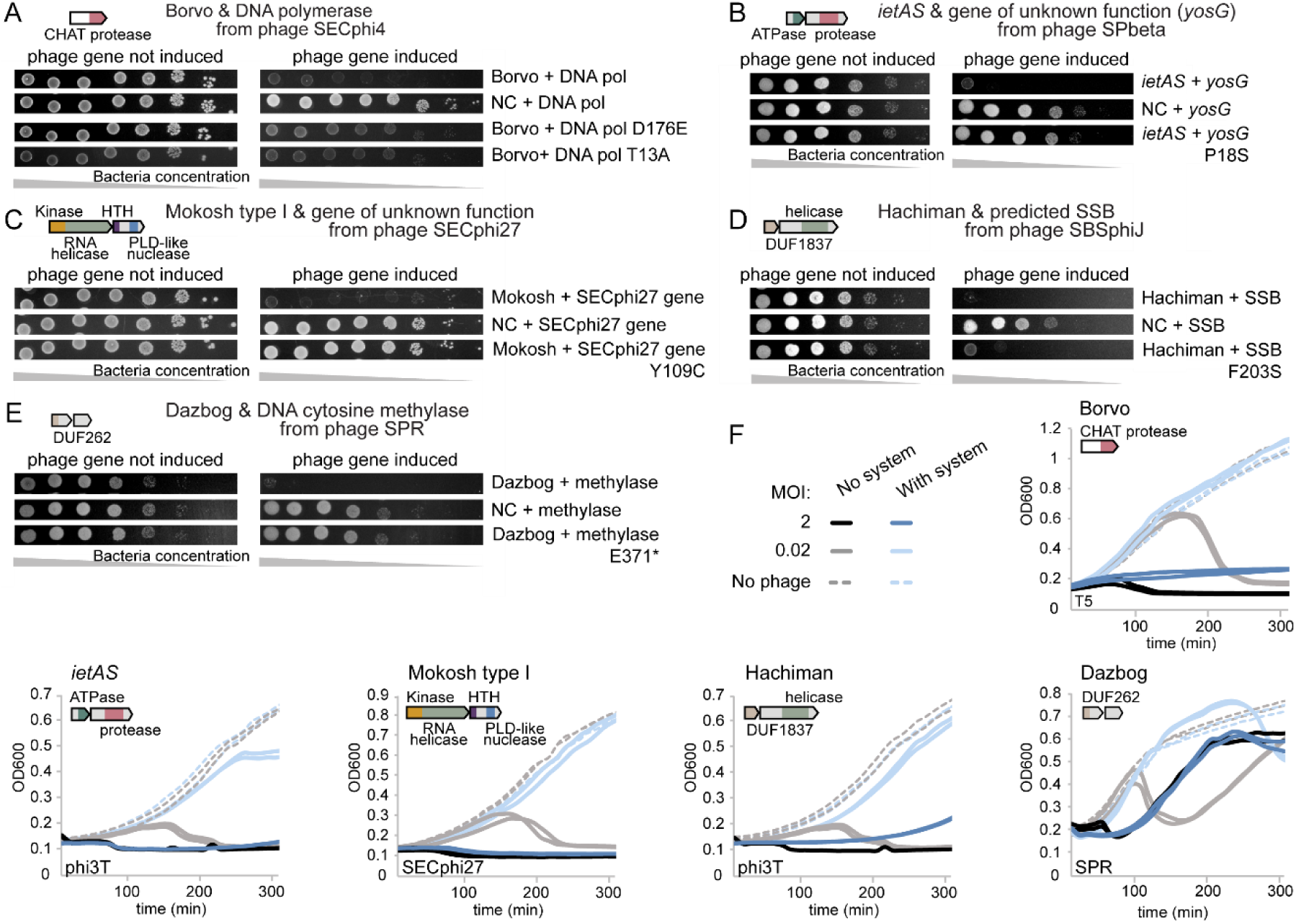
Identification of defense systems that function via abortive infection and the phage genes that activate them. (A-E) Expression of a single phage gene triggers toxicity in defense system-containing strains. Bacterial viability was measured by plating cells encoding the specified defense system and the phage gene, in ten-fold serial dilutions, under conditions that repress or induce expression of the phage gene. NC is a negative control strain lacking the defense system. Panels A-E show one representative of three replicates. (A) Expression of WT or mutated DNA polymerase from phage SECphi4 in *E. coli* MG1655 with the Borvo system from *E. coli* 403116. (B) Expression of WT or mutated *yosG* from phage SPbeta in *B. subtilis* BEST7003 with the *ietAS* system from *B. cereus* m1293. (C) Expression of WT or mutated protein of unknown function (128aa) from phage SECphi27 in *E. coli* MG1655 with the Mokosh type I system from *E. coli* ETEC H10407. (D) Expression of WT or mutated predicted ssDNA-binding protein (SSB) from phage SBSphiJ in *B. subtilis* BEST7003 with the Hachiman system from *B. cereus* B4087. (E) Expression of WT or mutated DNA cytosine methylase from phage SPR in *B. subtilis* BEST7003 with the Dazbog system from *B. mycoides GOE9*. (F) Growth curves in liquid culture for *E. coli* MG1655 or *B. subtilis* BEST7003 bacteria that contain the specified anti-phage system (“with system”) and bacteria that lack the anti-phage system (“no system”), infected with *E. coli* phage T5 or SECphi27 or *B. subtilis* phage phi3T or SPR. Bacteria were infected at time 0 at a multiplicity of infection (MOI) of 0.02 or 2, and the first measurement of the optical density (OD_600_) was taken 10 minutes post infection. Two biological replicates are presented for each experiment as individual curves. The two experiments with the phi3T phage were performed simultaneously with the same negative controls, and thus negative control curves are identical for the Hachiman and *ietAS* panels. The regrowth observed when the temperate phage SPR infects Dazbog-containing and Dazbog-lacking strains is attributed to the formation of SPR lysogens.

Although our approach reveals phage genes whose expression activates toxicity of abortive infection systems, it does not indicate whether the phage protein activates the defense system by direct binding or, alternatively, indirectly triggers the system. For example, the Dazbog system becomes toxic in cells that express a DNA cytosine methylase from phage SPR (Figure 5E). This methylase is known to methylate phage DNA at CCGG and other related motifs (Gunthert and Reiners, 1987). We found that the genome of SPR phages that escaped Dazbog became unmethylated at CCGG, showing that the escape mutation in the methylase abolished its DNA methylation function (Figure S4). It is therefore possible that Dazbog may be an abortive infection system triggered by the presence of methylated DNA, which is consistent with the growth arrest phenotype seen when Dazbog-expressing cells are infected at high MOI (Figure 5F). The first gene in the two-gene Dazbog system harbors a protein domain (DUF262, pfam accession PF03235) that is found in other defense systems including GmrSD and SspABCDE, which sense modified DNA (Liang et al., 2007; Machnicka et al., 2015; Xiong et al., 2020), supporting the hypothesis that Dazbog activity depends on the recognition of modified DNA.

### Recurring themes in phage escape from bacterial defense

The large number of escaper phages isolated in this study, and the richness of the defense system cohort studied reveal common recurring themes among the mutations that allow phage escape from bacterial defense. We identify three such main categories: components of the phage core replication machinery, phage structural proteins, and host takeover mechanisms.

#### Components of the phage core replication machinery

In many cases, escape mutations were detected in proteins involved in the core replication machinery of the phage, including the DNA polymerase, primase-helicase, and ssDNA-binding proteins (SSBs) (Figure 3, Figure 6). Such mutations allowed phages to escape the systems Lamassu, Hachiman, *ietAS*, Retron-Eco8, ShosTA, Borvo, and some AVAST systems (Figure 3, Figure 6). Previous studies on the anti-phage defense systems AbiK (Bouchard and Moineau, 2004; Wang et al., 2011), AbiQ (Samson et al., 2013b), ApbA/B (Yasui et al., 2014), DarTG (LeRoux et al., 2022), Tin (Mosig et al., 1997), and Nhi (Bari et al., 2022) have also reported phage escape via mutation in one of these core replication proteins. Thus, diverse defense systems have evolved to sense or target the core determinants of phage replication. As the phage replication machinery is essential for successful infection, it is advantageous for defense systems to converge on core components of this machinery as a trigger for infection recognition. Moreover, the essentiality of the replication complex would limit the possibility for loss-of-function escape mutations. Another advantage of sensing the SSB protein specifically, is that it is one of the most abundant proteins generated by the phage during infection (Hernandez and Richardson, 2019), offering ample targets for the defense systems to recognize.

**Figure 6.**
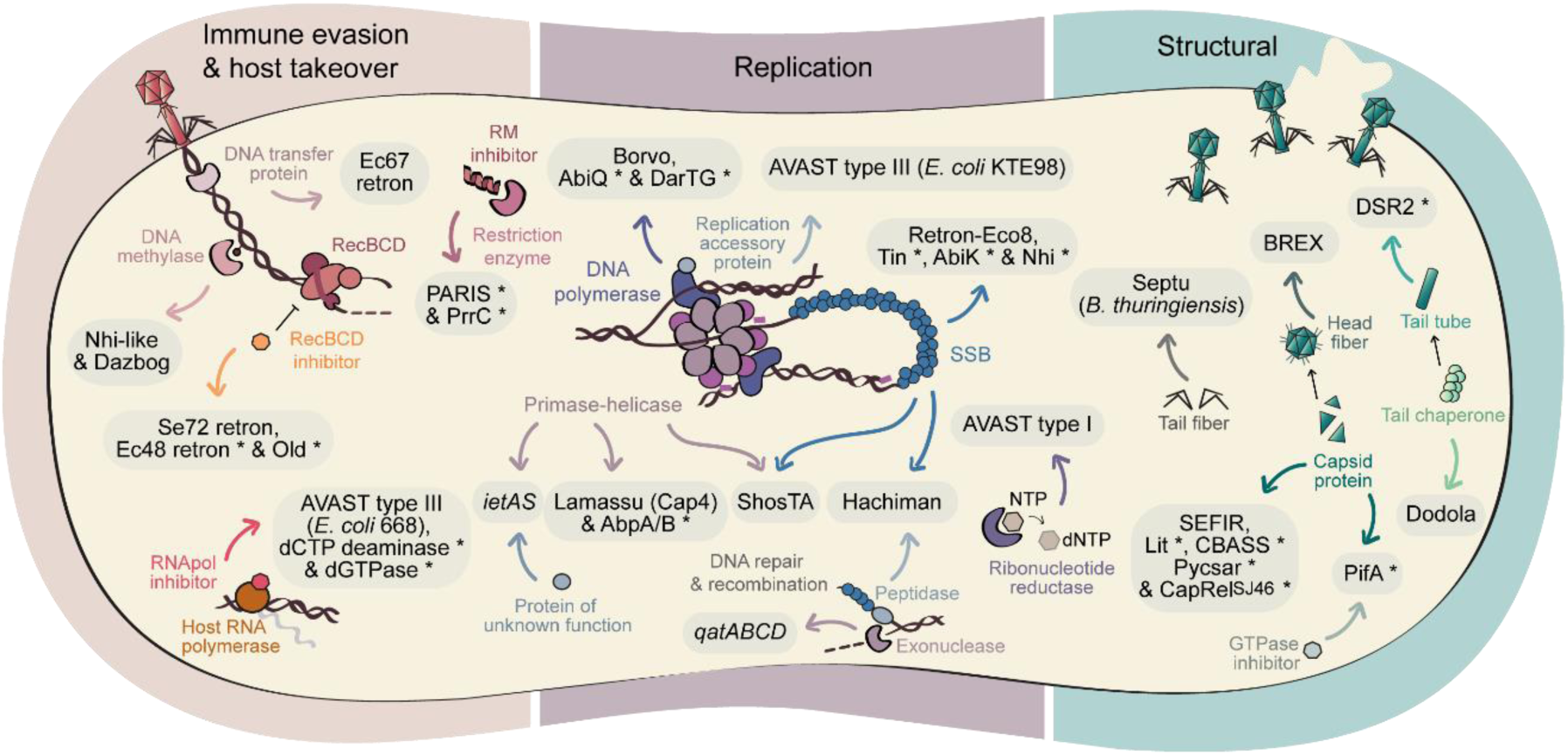
Diverse modes in which bacteria sense phage infection. Schematic depiction of the phage components that confer sensitivity to bacterial anti-phage systems listed in Figure 3B, as well as selected examples from previous studies. Proteins listed in Figure 3B that do not correspond to one of the three groups are not depicted. Phage sensitivity components identified in previous studies are marked here with an asterisk and are detailed in Supplementary Table S4.

In some cases, the same replication machinery gene was mutated in multiple phylogenetically distant phages to allow them to escape the same defense system. For example, the T5, SECphi4 and SECphi18 phages overcame the Borvo defense system via a missense mutation in the gene encoding the phage DNA polymerase (Figure 3B, Table S2). In all cases, escape mutations occurred in the proofreading cleft of the polymerase (Figure S5). Notably, two mutations were in residues that are highly conserved across DNA polymerase proteins (T13 and D176 in the SECphi18 DNA polymerase), and were shown to play an important role in the ssDNA binding and metal binding functions of the proofreading domain (Bernad et al., 1989; Kamtekar et al., 2004). As there is no clear sequence homology between the DNA polymerase proteins of T5 and the other two phages (SECphi4 and SECphi18), it is possible that this system senses the general structure of the polymerase, its interaction with other proteins, or the outcome of its interaction with DNA, as a signature for infection.

Similarly, phage escape from the Hachiman and ShosTA defense systems involved mutation in genes that participate in DNA replication, recombination and repair in various phages (for Hachiman, SSB proteins of the Gp2.5 and RecA families, and a DNA repair protein, and for ShosTA, SSB and Primase-Helicase proteins). The fact that mutations in different DNA replication and manipulation proteins enable phage escape from defense suggests that Hachiman and ShosTA may respond to a protein-DNA complex produced as an intermediate of phage DNA replication or recombination, rather than to a specific phage protein, as suggested for the RexAB abortive infection system (Snyder, 1995). Interestingly, the ShosTA system was previously shown to encode a toxin-antitoxin system (Kimelman et al., 2012), whereby the antitoxin shows homology to DprA, a protein that binds to ssDNA and SSB proteins (Mortier-Barrière et al., 2007; Rousset et al., 2022). It is therefore possible that the ShosTA DprA-like protein acts as the sensor of this system, triggered by a phage replication intermediate that involves ssDNA.

#### Phage structural proteins

A second major category of escape mutations involves mutations in phage structural genes (Figure 3, Figure 6). For example, mutations in the phage capsid protein enable various phages to overcome the defense systems SEFIR (Figure 3B, Table S2), Pycsar (Tal et al., 2021), some types of CBASS (Huiting et al., 2022), PifA (Molineux et al., 1989), Lit (Champness and Snyder, 1982), and CapRel^SJ46^ (Zhang et al., 2022). Most of these systems function via abortive infection, but at least in the case of CBASS, Pycsar and SEFIR, co-expression of the wild-type form of the major capsid protein with the defense system did not activate toxicity of the system (Figure S3A) (Huiting et al., 2022; Tal et al., 2021). This implies that the system might be triggered by a higher order structure of the capsid. Alternatively, the system could be activated by the interaction of the capsid protein with other phage or host factors, as shown for the Lit abortive infection protein which is triggered when the major head protein of phage T4 interacts with the translation elongation factor EF-Tu (Bingham et al., 2000). In other cases, direct interaction of the phage structural protein with the defense system results in activation of abortive infection, as recently demonstrated for the DSR2 (Garb et al., 2021), and CapRel^SJ46^ (Zhang et al., 2022) systems. We note that it makes biological sense for abortive infection systems to sense phage structural genes, which are typically expressed at late stages of the phage infection cycle and are usually highly expressed. In this way, an abortive infection system will elicit cell death or growth arrest as a last resort when recognizing that phage infection has advanced to its late stages.

Phages that escaped the *B. thuringiensis* Septu defense system had mutations in their tail fiber genes (Figure 3). Interestingly, this system provides defense against both *E. coli* and *B. subtilis* phages when expressed in either *E. coli* MG1655 *or B. subtilis* BEST7003, respectively. We isolated *E. coli* T7 phages and *B. subtilis* SBSphiJ phages that overcome Septu defense, and, although these phages are very distant from each other and infect completely different hosts, they both escaped the Septu system upon mutation in the phage tail fiber gene (Figure 3B). Moreover, for both the T7 and SBSphiJ phages, the mutations were located at the C-terminal tip of the tail fiber proteins, suggesting a similar recognition mechanism despite no detectable sequence similarity between the tail fibers of these phages (Table S2). On the other hand, escape from a different variant of the Septu defense system, cloned from *B. weihenstephanensis,* occurred upon mutation in different phage genes apparently unrelated to the phage tail fiber (Figure 3B), suggesting that different variants of the same system may sense different phage components.

#### Host takeover mechanisms

A third category of escape mutations occur in genes involved in host takeover and immune evasion. These include genes whose protein products inhibit RecBCD, thus activating retrons Se72 and Ec48 (Millman et al., 2020), as has also been shown for the P2-encoded protein Old (Lindahl et al., 1970; Myung and Calendar, 1995). Another example are proteins that inhibit restriction-modification systems, resulting in activation of the PrrC (Penner et al., 1995) and PARIS (Rousset et al., 2022) abortive infection systems. These systems are “guardians” of core defense machineries in the cell and become activated as a second line of defense if the phage tampers with the guarded system (Millman et al., 2020). Another example of how phage evasion of one defense system can serve as a trigger of another system is the case of the Dazbog and Nhi-like systems. In both cases, loss of function mutations in a phage DNA methylase enable the phage to overcome defense (Figure 3B, Figure S4). Thus, methylation of phage DNA, which normally protects phages from cleavage by restriction-modification systems, likely sensitizes them to the Dazbog and Nhi-like systems. Phage takeover and manipulation of the host transcription machinery also seems to be the focal point for sensing by several defense systems. These include the type III AVAST system from the *E. coli* 668 strain (Figure 3) and the defensive deoxynucleotide depletion enzymes dCTP deaminase and dGTPase (Tal et al., 2022); mutations in T7 phage genes responsible for inhibition of host transcription allow escape from these three systems. General inhibition of host transcription by phage was previously found to trigger also the toxin-antitoxin system *toxIN* (Guegler and Laub, 2021).

## Discussion

In this study, we attempted to address the key question of how bacterial immune systems sense phage infection. By screening for phage suppressor mutants that can overcome bacterial immunity, we identified the components sensitizing phages to 19 bacterial defense systems. While previous studies focused on individual systems, our study shows that there are unifying principles in phage sensing among highly diverse defense systems. We demonstrate that multiple systems have evolutionarily converged to sense the same phage component as a signal for infection, and found that the determinants sensitizing phages to bacterial immune systems fall mostly within three general categories: the core DNA replication machinery, phage structural proteins, and components involved in host immune evasion and takeover. While 75% of the genes in a typical phage encode “hypothetical proteins” of unknown function (Edwards and Rohwer, 2005), the majority of genes mutated in escaper phages were of known functions, suggesting that defense systems tend to target core rather than accessory components of the phage biology.

While the large number of phage mutants described here cannot be fully explored mechanistically in a single study, we were able to identify specific phage proteins necessary and sufficient for activation of multiple defense systems. These include the Borvo and Hachiman systems that sense phage DNA polymerase and SSB, respectively; the Dazbog system that responds to phage DNA methylase activity; the *ietAS* and Mokosh systems that are triggered by phage proteins of unknown function; and two retron systems, one responding to RecBCD inhibition by the λ Gam protein and the other activated by phage SSB. Although we show that expression of each individual phage protein triggers the respective defense system, our data does not reveal whether this triggering is by direct interactions of the phage protein and the defense system or by an indirect effect. Previous studies have identified defense systems that are activated by direct binding to a phage protein, as in the case of DSR2 (Garb et al., 2021) and CapRel (Zhang et al., 2022). On the other hand multiple defense systems are known to sense the effect of the phage protein and not the protein itself, for example retrons Ec48 and Se72 that sense RecBCD inhibition (Millman et al., 2020; this study), as well as PrrC that is triggered upon sensing the inhibition of a restriction enzyme by phages (Penner et al., 1995). Further studies would be necessary to elucidate how each of the phage components we identified directly or indirectly activates the respective systems.

Some of the phage mutants we isolated may not escape by mutating the component identified by the defense system, but rather evade immunity via other means. It is possible that some mutations could allow the phage to overcome the effect of the system, as proposed, for example, in the case of the DarTG system, which installs ADP-ribose on phage DNA in response to infection (LeRoux et al., 2022). Phages mutated in their DNA polymerase could tolerate the effect of the DarTG system, and it was suggested that these mutations allowed the polymerase to replicate DNA despite the DNA modifications made by the DarTG system (LeRoux et al., 2022). Another possibility is that the mutation in the phage has a “gain of function” effect, generating a protein that actively inhibits the defense system. Such mutants were described in two recent studies in which mutations switched a phage anti-CBASS or anti-DarTG protein from an inactive to an active form, enabling phages to overcome the CBASS and DarTG systems, respectively (Huiting et al., 2022; LeRoux et al., 2022). Another study has identified phage mutations that allowed phages to overcome the ToxIN system by evolving a non-coding RNA acting as an antitoxin to block the ToxN protein (Blower et al., 2012).

Notably, despite our success in isolating escape mutants for many systems, such mutants may not easily arise in nature. Previous studies have shown that mutant phages escaping from defense systems have fitness defects, and become extinct in competition assays with WT phages when infecting bacteria without the defense system (Tal et al., 2021). In agreement with this, we observe that the mutant phages escaping defense in our study often formed smaller plaques on control cells lacking the defense system as compared to the WT phage, suggesting that the mutation enabling phage escape comes with a fitness cost. Moreover, most bacteria encode more than one defense system, with an average of ∼5 systems per genome (Millman et al., 2022; Tesson et al., 2022). Thus, in natural settings, we expect it to be more difficult for phages to escape bacterial immunity by a single mutation. Indeed, in some cases defense systems have been shown to act together to enhance anti-phage protection and limit phage escape (Dupuis et al., 2013; Maguin et al., 2022; Silas et al., 2017).

For 35 systems included in our initial screen we were unable to isolate phages that escaped defense. This highlights the limitations in our experiment, and suggests there may be additional immune sensing strategies undetected by our method. In these cases, it is possible that the phage component sensed by the system is essential, such that mutations that escape detection by the system would also render the phage inactive. It is also possible that escape may require more than one mutation event in the phage, such that the probability to achieve a combination of these mutations was too low in our experimental setup.

The plethora of bacterial anti-phage immune systems that were recently discovered are still largely unexplored mechanistically. In addition to the direct insights derived from our study, we envision that our data and the large number of phage mutants we have collected will serve as a valuable resource for future studies aiming to solve the mechanisms of multiple defense systems. We also foresee that the general principles for phage sensing we describe in this study will hold true for other defense systems discovered and studied in the future.

## Materials and Methods

### Bacterial strains and phages

Defense systems were expressed in either *E. coli* MG1655 (ATCC 47076) or *B. subtilis* BEST7003 (obtained from M. Itaya at Keio University, Japan). *E. coli* BW25113 knockout strains (Δ*recB* and Δ*iscR*) from the Keio collection (Baba et al., 2006) were obtained from the Weizmann institute bacteriology repository (#JW2788-1 (*RecB*), #JW2515-3 (*iscR*)). Bacteria were grown in MMB (LB supplemented with 0.1 mM MnCl_2_ and 5 mM MgCl_2_) at 37°C shaking at 200 r.p.m, unless specified otherwise, and the appropriate antibiotics were added. For genes expressed in *E. coli*, ampicillin (100 μg/ml) or kanamycin (50 μg/ml)) was used to ensure maintenance of plasmids. For genes expressed in *B. subtilis*, spectinomycin (100 μg/ml) or chloramphenicol (5 μg/ml) was used to ensure the presence of an integrated antibiotics resistance cassette in the *B. subtilis* genome.

The phages used in this study are listed in Supplementary Table S5. Infection was performed in MMB with or without agar (0.3% or 0.5% as detailed in Supplementary Table ^S5^). For all *Bacillus* phages, MMB was supplemented with 5 mM CaCl_2_ for all plaque assays.

### Plasmid and strain construction

Defense systems included in this study are listed in Table S1. For defense system-expressing strains that were constructed previously, the reference study describing the details regarding the construction of the strain is provided (Table S1). For the defense systems reported by Gao et al. including RADAR, AVAST, qatABCD, Ppl, TerY-p and *ietAS*, a homolog of the system from the specified donor organism (Table S1) was cloned into the pSG1 shuttle vector (Doron et al., 2018) together with their native promoters between the AscI and NotI sites of the multiple cloning site. The sequences of these systems are provided in Table S1. DNA sequences of the RADAR, AVAST, *qatABCD*, Ppl and TerY-p systems were synthesized and cloned by Genscript Corp. The *ietAS* system was amplified from genomic DNA of the donor strain using KAPA HiFi HotStart ReadyMix (Kapa Biosystems KK2601) with primers OSM646 and OSM647 (Table S6), and cloned using the NEBuilder HiFI DNA Assembly cloning kit (NEB E5520S) into the pSG1 vector (amplified with primers OSM315 and OSM316 (Table S6)).

Three of the systems were cloned by Doron et al. (2018) for expression in *B. subtilis* BEST7003 (Lamassu from *Bacillus* sp. NIO-1130, Septu from *B. thuringiensis*, and Thoeris from *B. amyloliquefaciens*). In this study, we used the pSG1 shuttle vectors expressing these three systems constructed by Doron et al. (2018) for expression of the systems in *E. coli* MG1655 as well. Expression of these systems in *E. coli* provided defense against several different *E. coli* phages, which were therefore included in our screen (Figure 2).

Defense system containing vectors were transformed into either electrocompetent *E. coli* MG1655 or using MC medium into *B. subtilis* BEST7003 (Wilson and Bott, 1968). From an overnight culture of *B. subtilis*, 60 μl were diluted in 2700 μl ultra-pure water, 30 μl 1M MgSO_4_ and 300 μl 10X MC medium (80 mM K_2_HPO_4_, 2% glucose, 30 mM trisodium citrate, 22 μg/ml ferric ammonium citrate, 0.2% sodium glutamate, 30 mM KH_2_PO_4_ and 0.1% casein hydrolysate (Merck Millipore 102245)). The bacteria were incubated at 37°C, 200 r.p.m until reaching an OD_600_ of 0.6. Then 300 μl was transferred to a new 15 ml tube and 300 ng of plasmid DNA was added. The tube was incubated for another 3 hours (37°C, 200 r.p.m), and then plated on LB agar plates supplemented with 100 μg/ml spectinomycin and incubated overnight at 30°C. The resulting transformants were screened on starch plates for an amylase-deficient phenotype to identify integration of the defense system into the *Bacillus amyE* locus.

As negative control strains lacking an inserted defense system, an empty pSG1 plasmid was used without any system inserted in the multiple cloning site. For the *E. coli* negative control (NC) strain, the empty plasmid was transformed into *E. coli* MG1655. For the *Bacillus* NC strain, the empty plasmid was transformed into *B. subtilis* BEST7003 and the spectinomycin-resistance gene was integrated into the *amyE* locus.

The majority of the phage genes expressed in this study were synthesized and cloned by Genscript Corp. A list of all phage genes expressed in this study is provided in Table S3. For expression in *E.* coli, the phage gene was cloned under an arabinose-inducible P_BAD_ promoter, by replacing the RFP ORF in either pBbS8k-RFP or pBbE8k-RFP (Addgene Plasmids #35276 and #35270). For expression in *B. subtilis*, the phage gene was cloned under an IPTG inducible promoter (P_hy-spank_) by replacing the sfGFP ORF in the *Bacillus* integration vector pSG-*thrC*-P_hy-spank_-sfGFP (Garb et al., 2021) or pDR111-sfGFP (Overkamp et al., 2013) for genomic integration in the *Bacillus thrC* or *amyE* locus, respectively. The SECphi4 56aa protein of unknown function was the only gene not synthesized by Genscript Corp. The gene encoding this protein was amplified from genomic DNA of the SECphi4 phage using KAPA HiFi HotStart ReadyMix (Kapa Biosystems KK2601) with primers AS330+AS331 (Table S6), and cloned using the NEBuilder HiFI DNA Assembly kit into the pBbE8k-RFP vector (amplified with primers AS328+AS329 (Table S6)).

Vectors expressing each of the phage genes were transformed into defense system-containing cells or negative control cells lacking the system (for all cases besides the predicted tail fiber gene of phage SBSphiJ, see note below). Transformation to *E. coli* strains expressing defense systems was performed using either chemical transformation in TSS medium (LB supplemented with 10% (w/v) PEG 8000, 0.6% (w/v) MgCl_2_*6H_2_0, and 5% (v/v) DMSO) (Chung and Miller, 1993) or by electroporation into electrocompetent cells expressing the required defense system. Following transformation, the cells were recovered in LB media supplemented with 1% glucose to repress leaky expression from the P_BAD_ promoter, and then plated on LB agar plates supplemented with 50 μg/ml kanamycin + 1% glucose. Transformation to *Bacillus* strains expressing defense systems was performed using MC medium as described above, and incubated overnight at 30°C on LB agar plates supplemented with 5 μg/ml chloramphenicol or 100 μg/ml spectinomycin. For expression of the SBSphiJ predicted tail fiber gene, we found that the *B. thuringiensis* HD12 Septu defense system inhibited transformation when expressed in *B. subtilis* BEST7003, and we therefore first cloned the phage gene into the bacterial genome at the *thrC* locus, and then introduced the Septu defense system into the *amyE* locus.

### Isolation of mutant phages that overcome defense

To screen for mutant phages that escape defense, phages were plated on bacteria expressing each of the defense systems (Table S1) using the double-layer plaque assay (Kropinski et al., 2009) (Figure 1A). For this, 100 μl of bacterial cells grown in MMB to an OD_600_ of 0.3 were mixed with 100 μl phage lysate. After 10 minutes at room temperature, 5 ml pre-melted 0.3% or 0.5% MMB agar was added (Table S5), and the mixture was poured onto MMB 1.1% agar plates. The double layer plates were incubated overnight at 37°C or room temperature (Table S2) and single plaques were picked into 90 μl phage buffer (50 mM Tris pH 7.4, 100 mM MgCl2, 10 mM NaCl)). In addition to picking single plaques, the entire top layer was scraped into 2 ml phage buffer to enrich for phages that escape defense. Phages were left for 1 hour at room temperature during which the phages were mixed several times by vortex to release them from the agar into the phage buffer. The phages were centrifuged at 3200 g for 10 min to remove agar and bacterial cells, and the supernatant was transferred to a new tube.

In order to test the collected phages for their ability to escape from the defense system they formed plaques on, the small drop plaque assay was used (Mazzocco et al., 2009) (Figure 1B). 300 μl of an overnight culture of bacteria expressing each defense system or a negative control strain lacking the defense system, were mixed with 30 ml melted 0.3% or 0.5% MMB agar (Table S5). The mixture was poured onto a 10-cm square plate and left to dry for 1 hour at room temperature. Ten-fold serial dilutions in phage buffer were performed for the ancestor phages (WT phage used for the original double layer plaque assay) and the phages collected from plaques formed on the defense system strain. 10 μl drops of each phage were plated on the bacterial layer. The plates were incubated overnight at 37°C or room temperature (Table S2). Plaque-forming units (PFUs) observed after overnight incubation were counted and used to calculate the fold defense using the ratio between phage PFUs obtained on defense-containing cells and PFUs obtained on negative control cells. When individual plaques could not be deciphered, a faint lysis zone across the drop area was considered to be 10 plaques.

### Amplification of mutant phages

Isolated phages for which there was decreased defense compared to the ancestor phage were propagated to obtain a high titter phage stock for sequencing. For this, a single plaque formed on the defense strain in the small drop plaque assay was picked into a liquid culture of defense system expressing cells grown in 1 ml MMB to an OD_600_ of 0.3. The phages were incubated with the bacteria at 37°C 200 r.p.m shaking for 3 hours, and then an additional 9 ml of bacterial culture grown to OD_600_ 0.3 in MMB was added, and incubated for another 3 hours (37°C 200 r.p.m). The lysate was then centrifuged at 3200 g for 10 min and the supernatant was filtered through a 0.2 μM filter to get rid of remaining bacteria. Phage titer was then measured using the small drop plaque assay on the negative control strain and in cases where the titer was less than 10^7^ PFU/ml, the phage titer was raised using either another round of propagation in liquid culture or with a plate lysate (Kropinski et al., 2009).

For plate lysates, 10^3^-10^5^ PFUs formed on defense system expressing cells using a double-layer plaque assay were scraped into 5 ml of phage buffer. After 1 hour at room temperature, the phages were centrifuged at 3200 g for 10 min, and the supernatant was filtered through a 0.2 mM filter.

### Sequencing and genome analysis of phage mutants

DNA was extracted from 500 μl of a high titer phage lysate (> 10^7^ PFU/ml). The phage lysate was treated with DNase-I (Merck cat #11284932001) added to a final concentration of 20 μg/ml and incubated at 37°C for 1 hour to remove bacterial DNA. DNA was then extracted using the QIAGEN DNeasy blood and tissue kit (cat #69504) starting from the Proteinase-K treatment step to lyse the phages. Libraries were prepared for Illumina sequencing using a modified Nextera protocol (Baym et al., 2015). Reads were aligned to the phage reference genomes (NCBI accession numbers provided in Table S5) and mutations compared to the reference genome were identified using Breseq (version 0.29.0) with default parameters (Deatherage and Barrick, 2014). Only mutations that occurred in the isolated mutants, but not in the ancestor phage, were considered. Silent mutations within protein coding regions were disregarded as well. Mutations marked by Breseq under “marginal evidence” were reported if the mutation was supported by more than 20% of the aligned reads, and the percent of the mutated reads for such cases is provided in Table S2.

### Bacterial growth upon induction of phage genes

For each of the defense systems, at least one phage sensitivity component identified in our screen was chosen for cloning. A plasmid containing each of the phage genes under an inducible promoter (P_hy-spank_ or P_BAD_) (Table S3) was transformed into defense system-containing bacteria or negative control cells lacking the defense system as explained above. After overnight incubation, 8 to 16 of the freshly transformed colonies were then streaked on LB agar plates with or without added inducer (1mM IPTG or 0.3% arabinose (Table S3)). Bacterial growth was examined after overnight incubation at 37°C for *E. coli* or 30°C for *Bacillus*.

The toxicity of the phage genes was measured quantitatively by counting the number of bacterial colony-forming units (CFUs) per ml formed after inducing expression of the phage gene overnight. For genes expressed in *Bacillus*, two loopfuls of bacteria were taken from 3 different streaked colonies that grew without added inducer and inoculated directly in 90 μl phosphate-buffered saline (PBS) (Thermo Fisher Scientific 14200075). For genes expressed in *E. coli*, 3 different streaked colonies that grew without added inducer were inoculated in MMB supplemented with 100 μg/ml ampicillin + 50 μg/ml kanamycin + 1% glucose and grown to stationary phase at 37°C 200 r.p.m. Then, ten-fold serial dilutions of the bacteria in PBS were performed and 5 μl drops were spotted on LB agar plates with or without added inducer, either IPTG or arabinose, for expression of the phage gene. Inducer levels used for each gene are detailed in Table S5. Since some phage genes were toxic upon overexpression at maximal inducer levels (1mM IPTG or 0.3% arabinose), the level of inducer was lowered to reach a point where expression was not toxic in control cells. Plates were incubated overnight at 37°C for *E. coli* or 30°C for *Bacillus*, and bacterial colonies imaged and counted.

For both bacterial colony streaking and CFU experiments, non-induced plates were supplemented with 1% glucose to inhibit leaky expression of phage genes expressed under the P_BAD_ promoter.

### Transformation efficiency

Strains from the *E. coli* Keio collection (Δ*recB* and Δ*iscR* as a negative control) were used as recipients for chemical transformation in TSS media (Chung and Miller, 1993). An overnight culture of the recipient bacteria grown in MMB with 50 μg/ml kanamycin was diluted 1:100 into 5 ml of fresh media and grown at 37°C 200 r.p.m to OD_600_=0.2. Bacteria were then centrifuged at 3200 g for 10 minutes, after which the bacterial pellet was resuspended in 220 μl cold TSS media and put on ice. 50 μl of the bacteria were transferred to a new Eppendorf tube, and 100 ng of the vector expressing a retron defense system (pEc48, pSe72 or pEco8) or a negative control empty vector (pNC) was added. The bacteria were incubated with the vector on ice for 5 minutes, then at room temperature for 5 minutes, and again on ice for 5 minutes. 950 μl MMB was added and the bacteria were incubated at 37°C 200 r.p.m for 1 hour. Ten-fold serial dilutions of the bacteria in PBS were then plated on LB agar plates supplemented with 100 μg/ml ampicillin. Transformation efficiency was assessed by counting single colonies formed after overnight incubation at 37°C.

### Phage-infection dynamics in liquid medium

Overnight cultures of bacteria (*E. coli* MG1655 or *B. subtilis* BEST7003 expressing a defense system (Table S1)) or a negative control strain (containing the empty pSG1 vector) were diluted 1:100 in MMB medium supplemented with either ampicillin (100 μg/ml) for *E. coli* strains or spectinomycin (100 μg/ml) for *B. subtilis* strains. The diluted cultures were incubated at 37 °C 200 r.p.m for 100 min, after which 180 μl of the culture was transferred into a 96-well plate containing 20 μl of either phage buffer (for uninfected control) or phage lysate for a final MOI of 2 or 0.02. Plates were incubated at either 37 °C (for phage T5), 25°C (for phages SECphi27 and phi3T) or 30°C (for phage SPR) with shaking in a TECAN Infinite200 plate reader and OD_600_ was measured every 10 min. Infections were performed in duplicate from overnight cultures prepared from two separate colonies.

### Phage DNA restriction assay

High concentration phage DNA was prepared by first precipitating phages from a large volume, followed by DNA extraction using phenol-chloroform-isoamylalcohol. For this, 20 ml phage lysate was mixed with 4 ml 5M NaCl and 2 gr PEG8000. The samples were left to rotate overnight at 4°C, and then centrifuged at 11,000 g for 10 min at 4°C. The supernatant was discarded and the pellet was resuspended in 400 μl phage buffer (5mM Tris pH 7.4, 100 mM MgCl2, 10 mM NaCl). The samples were then treated with DNase-I (Merck cat #11284932001) added to a final concentration of 40 μg/ml and incubated at 37°C for 1 hour to remove bacterial DNA. Phages were then lysed by the addition of 200 μg/ml Proteinase-K (Bio-Lab cat #1673238300), 0.5% SDS (Bio-Lab cat #19812323) and 20 mM EDTA, incubated at 55°C for 1 hour. The lysate was then mixed with 1 volume of phenol-chloroform-isoamylalcohol (Merck cat #P2069), and centrifuged for at maximal speed in a table top centrifuge at 4°C for 15 min. The upper aqueous fraction was transferred to a new tube. To precipitate the DNA, 1 volume of isopropanol and 1/30 of the volume of NaAc 3M pH 6.5 were added to the collected aqueous fraction and mixed by inversion followed by incubation at −80°C for 25 min. Then the samples were centrifuged at maximal speed in a table top centrifuge at 4°C for 30 min, and the pellet was washed twice with cold 75% ethanol. The pellets were dried at 37°C for 5 min and resuspended in 50 μl ultra-pure water. DNA concentration was measured using a Qubit dsDNA BR Assay Kit (Thermo Fisher cat # Q32850). For the restriction assay, 300 ng phage DNA was mixed with 14 μl lysis buffer (50 mM Tris pH 7.5, 150 mM KCl, 1mM MgCl2, 1mM MnCl2, 1mM CaCl2) and 1 μl HpaII restriction enzyme (Thermo Fisher cat #ER0511). The reaction was incubated for 2 hours at 37°C and then loaded into a 0.7% agarose gel for analysis of cleavage pattern.

### Structure prediction and alignment of phage proteins

The structures for the SECphi4 phage putative SSB and T5 phage A1 proteins (GenBank accessions QJI52528.1 and AAS77051.1, respectively) were predicted using AlphaFold2 (Jumper et al., 2021) within ColabFold (Mirdita et al., 2022) with default parameters. For the SECphi4 SSB protein, the best ranking AlphaFold2 predicted structure was aligned to the T7 phage SSB (PDB:1JE5, chain A) using the “cealign” method of the alignment plugin within the PyMOL program v. 2.5.1 (Schrödinger, 2015).

For the T5 phage A1 protein, the amino acid sequence was first submitted to HHpred (Zimmermann et al., 2018), which revealed a putative nuclease domain at the C-terminus of the A1 protein but no known domain at the A1 N-terminus. The predicted structure of the A1 N-terminal domain (residues 1-60) was extracted from the best ranking AlphaFold2 predicted structure of the protein and submitted to PDBeFold (Krissinel and Henrick, 2004), a protein structure comparison web service to identify structural homologs, retrieving a high confidence hit to the DNA-binding domains of the *Caulobacter crescentus* GcrA (PDB 5yiv, chain D). To visualize the alignment, the crystal structure of the GcrA DNA-binding domain was aligned to the AlphaFold2 predicted structure of the T5 phage A1 protein using the alignment plugin within the PyMOL program with the “cealign” method.

The structures for the SECphi4 and T5 phage DNA polymerase proteins (GenBank accessions QJI52526.1 and AAS77168.1, respectively) were predicted using AlphaFold2 (Jumper et al., 2021) within ColabFold (Mirdita et al., 2022) with default parameters, and the best ranking AlphaFold2 predicted structure out of five predictions was chosen. The domains of the proteins were predicted using HHpred (Zimmermann et al., 2018) (3’-5’ exonuclease domain, pfam accession PF01612.23, and polymerase domain, pfam accession PF00476.23). The putative location of the active site was marked by adding the predicted metals to the AlphaFold2 structure. The alphafold2 predicted structures were aligned to the crystal structure of the T7 DNA polymerase (PDB: 1T7P) (Doublié et al., 1998) using the alignment plugin within the PyMOL program with the “cealign” method, and the location of the metals in the T7 DNA polymerase active sites were overlaid on the SECphi4 and T5 predicted DNA polymerase structures to identify the active site pockets.

### Sequence alignment of retron RTs

The protein sequences of the Ec48 and Se72 retron RTs (Gene IDs in the IMG database (Chen et al., 2021): 2642317602 and 2633939248, respectively) were aligned to each other using BLASTp on the NCBI BLAST web interface (Johnson et al., 2008) with default parameters.

## Acknowledgements

We thank the Sorek lab members and A. Bernheim for fruitful discussion and comments on earlier versions of this manuscript. A.M. was supported by a fellowship from the Ariane de Rothschild Women Doctoral Program and, in part, by the Israeli Council for Higher Education via the Weizmann Data Science Research Center. R.S. was supported, in part, by the European Research Council (grant ERC-AdG GA 101018520), Israel Science Foundation (grant ISF 296/21), the Deutsche Forschungsgemeinschaft (SPP 2330, grant 464312965), the Ernest and Bonnie Beutler Research Program of Excellence in Genomic Medicine, the Minerva Foundation with funding from the Federal German Ministry for Education and Research, and the Knell Family Center for Microbiology.

## Supplementary Material

**Figure S1.**
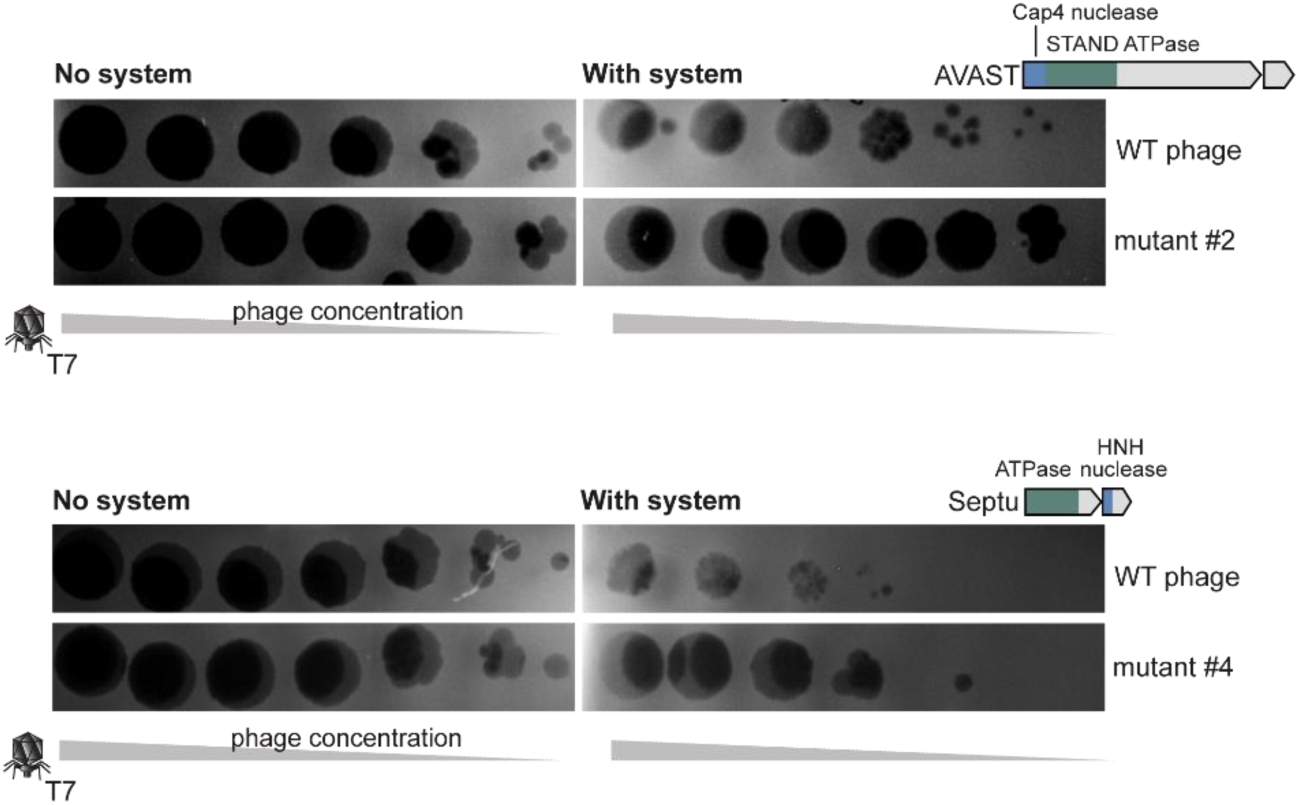
Escaper phages overcome reduction in plaque size. Several examples of cases where protection against the WT phage manifests in decreased plaque size and the escaping phages prevail, forming large plaques even when infecting bacteria with the defense system. Serial dilution plaque assays are presented for WT and mutant T7 phages infecting *E. coli* MG1655 expressing a defense system (AVAST type III from *E.coli* KTE98, or Septu from *B. thuringiensis* HD12).

**Figure S2.**
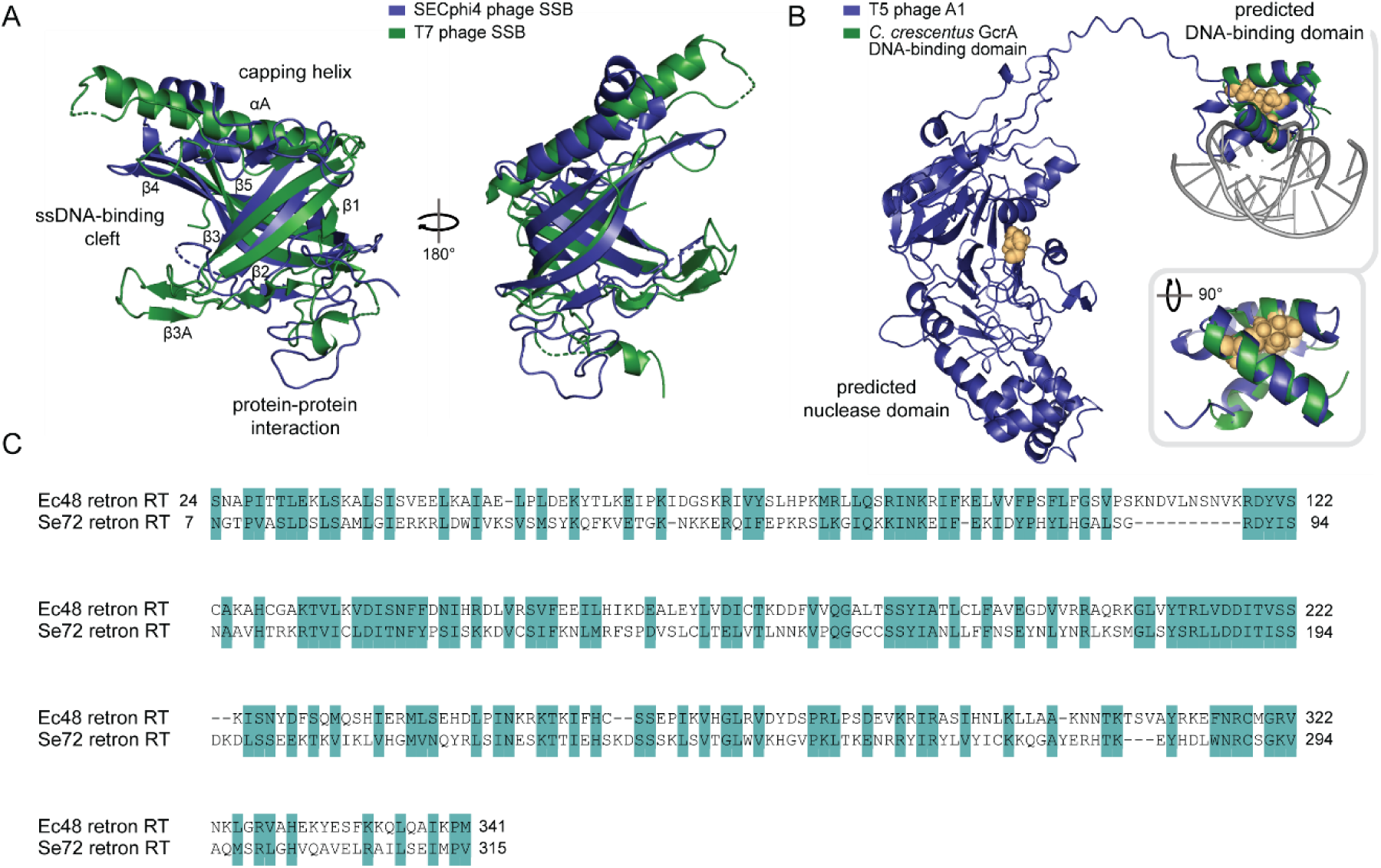
Analysis of components mutated in phages that escape retron defense systems. (A) Structural homology between the T7 phage single-stranded DNA-binding protein (SSB) and a predicted SSB from phage SECphi4. The AlphaFold2-predicted protein structure of the SECphi4 phage putative SSB (residues 1-186) is presented in blue, and the crystal structure of the T7 phage SSB (PDB: 1JE5, residues 1-196) is in green. Flexible C-terminal residues are not displayed for both proteins. Secondary structure elements of the ssDNA binding OB-fold of the T7 SSB are annotated based on (Hollis et al., 2001). (B) AlphaFold2-predicted protein structure of phage T5 A1 protein (blue). The N-terminal domain (residues 1-60) of the predicted A1 protein structure show structural homology to the *Caulobacter crescentus* GcrA DNA-binding domain (PDB 5yiv, chain D (green), bound to dsDNA (grey)). Amino acid residues mutated in T5 phages that overcome Ec67 retron-mediated defense are displayed as yellow spheres. (C) Amino acid sequence homology between the reverse transcriptase (RT) proteins of retron Ec48 and retron Se72. Identical and similar amino acids are shaded in turquoise.

**Figure S3.**
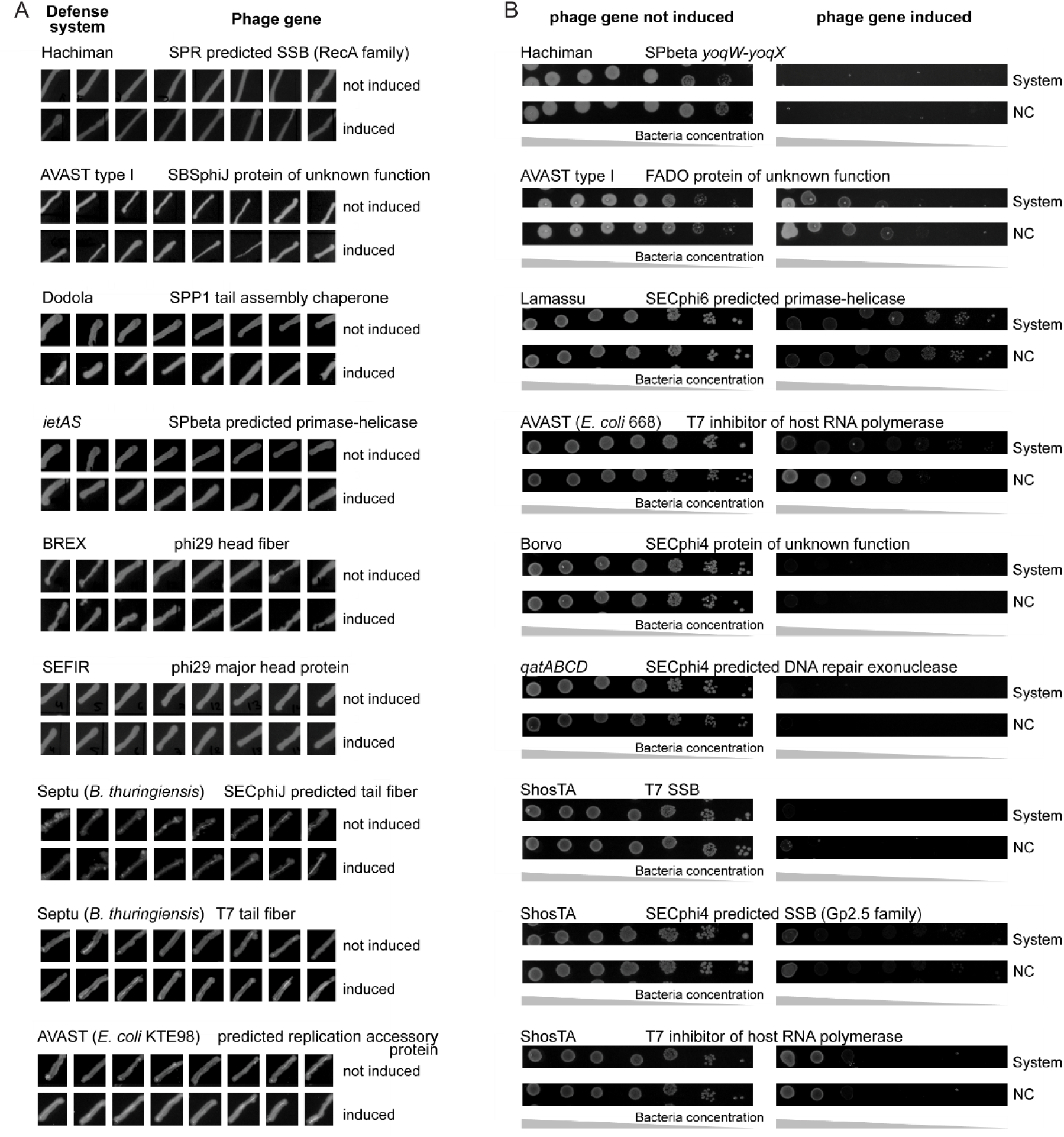
Expression of single phage genes in defense system-containing strains to identify defense systems that act via abortive infection and the phage protein that activates them. Bacterial viability was measured for cells encoding a defense system upon expression of single phage genes presented in Figure 3B. Bacterial growth was assessed by plating bacteria in conditions that repress or induce expression of the phage gene (with either arabinose or IPTG (see methods)). (A) Cases where expression of the phage gene was not toxic in cells that contained the defense system. Eight different bacterial colonies were streaked to assess toxicity upon expression of the phage gene in defense system-containing bacteria. (B) Cases where expression of the phage gene exhibited non-specific toxicity in both defense system-containing cells and in control cells lacking the system (NC). Measurement of the toxicity of the phage gene was performed by plating ten-fold serial dilutions of the bacteria. Images are representatives of three replicates.

**Figure S4.**
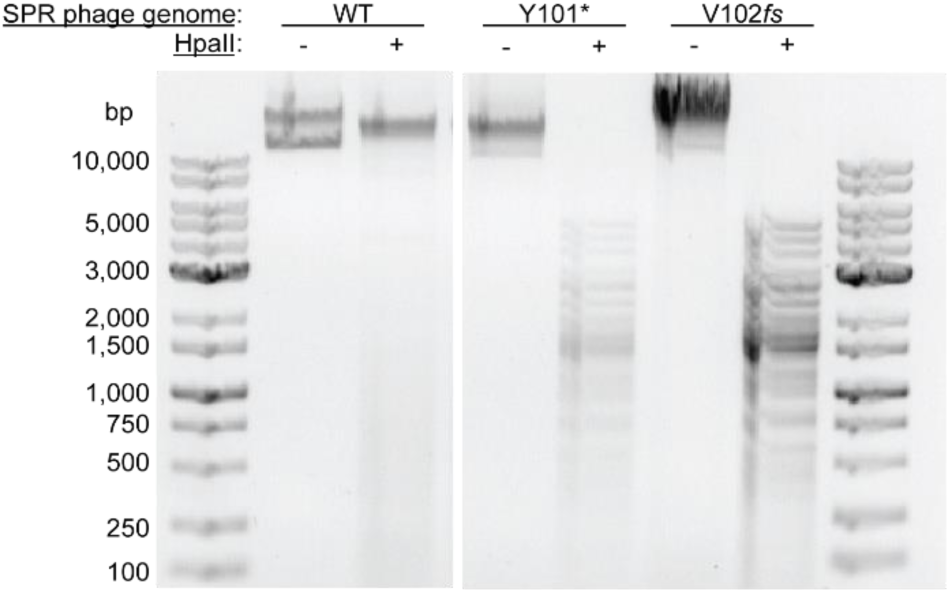
The genome of SPR escaper phages that overcome the Dazbog anti-phage system are unmethylated at CCGG. Restriction analysis of WT or mutated SPR phage DNA incubated for two hours with the HpaII restriction enzyme, which cleaves unmethylated DNA at CCGG motifs. Y101* and V102*fs* are mutations in the SPR DNA cytosine methylase that are found in phages that overcome Dazbog defense (Table S2). DNA bands were resolved by agarose gel electrophoresis in a 0.7% agarose gel. The genome of WT SPR phages is methylated at CCGG and is thus protected from cleavage by HpaII, while the genomes of mutant phages that escape Dazbog defense are restricted.

**Figure S5.**
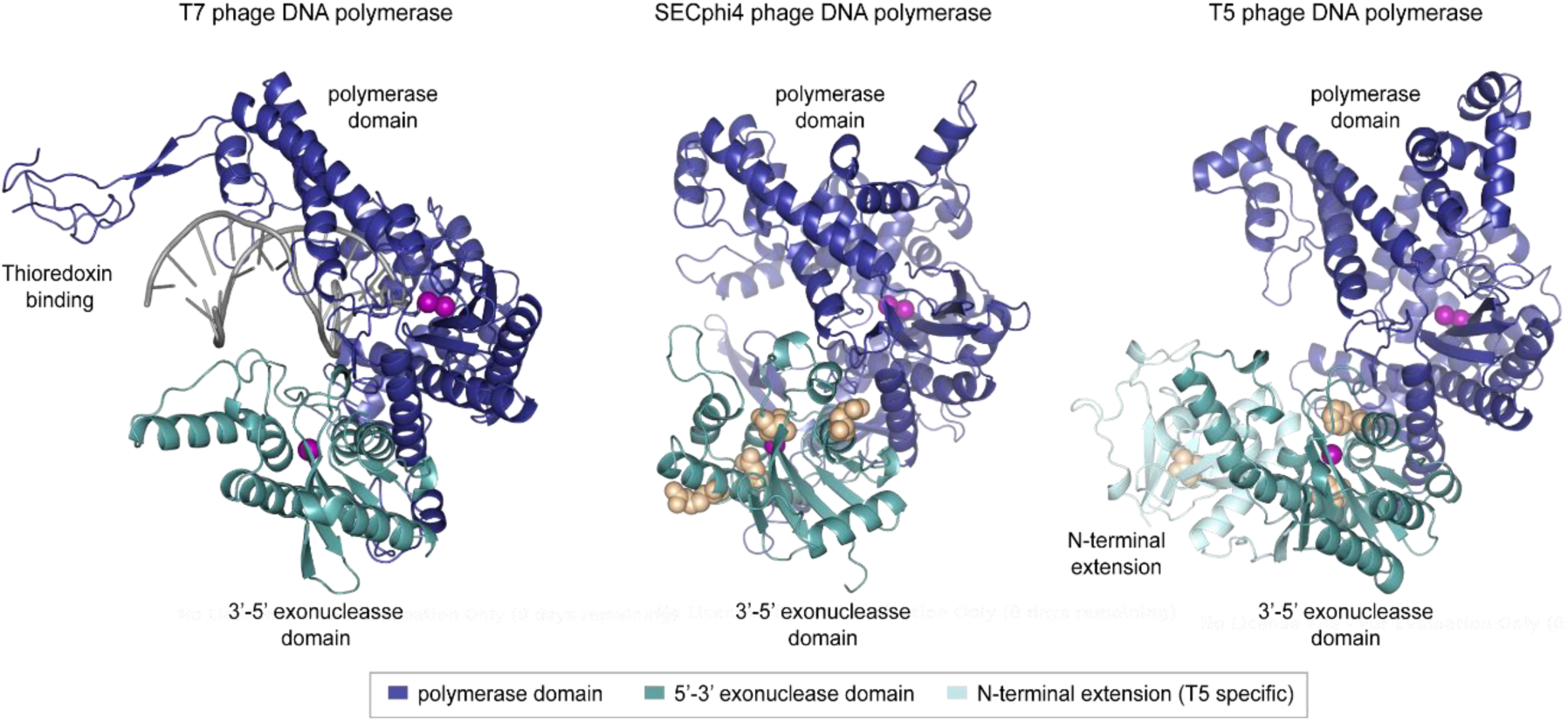
Mutations in the proofreading domain of phage DNA polymerase proteins enable escape from Borvo defense. AlphaFold2-predicted protein structures of the T5 and SECphi4 phage DNA polymerases are presented, as well as the crystal structure of the T7 phage DNA polymerase (PDB: 1T7P) (Doublié et al., 1998). Amino acid residues mutated in the DNA polymerase protein of phages T5 (I63, F223, T320), SECphi4 (T13, E91, D176) and SECphi18 (R171) that overcome Borvo-mediated defense are displayed as yellow spheres. The SECphi18 phage DNA polymerase is highly similar to that of SECphi4, thus SECphi18 mutations are marked on the SECphi4 DNA polymerase structure. The polymerase and proofreading (3’-5’ exonuclease) domains are colored in blue and cyan, respectively. T5 phage DNA polymerase contains an N-terminal extension (marked in light blue) that is absent in other DNA polymerases from the pol I family. This N-terminal extension is not part of the conserved 3’-5’ exonuclease, but might play a role in proofreading activity in T5, since its truncation reduces exonuclease activity (Andraos et al., 2004). Metals found in the active site of the polymerase and exonuclease domains are marked as magenta spheres based on their location in the T7 DNA polymerase crystal structure and are mapped onto the SECphi4 and T5 DNA polymerase predicted structures to mark their corresponding active sites.

**Table S1.** List of defense systems included in the screen and the source organism the system was taken from and cloned for heterologous expression in *E. coli* or *B. subtilis*. A reference is provided for systems cloned in other studies and for systems cloned in this study, the cloned sequence is provided.

**Table S2.** Mutations found in phages that escape defense

**Table S3.** List of phage genes cloned in this study

**Table S4.** List of previously reported genes mutated in phages that overcome defense

**Table S5.** Phages used in this study

**Table S6.** Primers used in this study

